# Characterising the balance between short-interval intracortical inhibition and short-interval intracortical facilitation in younger and older adults

**DOI:** 10.1101/643841

**Authors:** B.K. Rurak, G.R Hammond, H. Fujiyama, A.M. Vallence

**Author notes:** Correspondence to: Ann-Maree Vallence, Address: Discipline of Psychology, College of Science, Health, Engineering and Education, Murdoch University, 90 South Street, Murdoch WA 6150.

## Abstract

Measures of short-interval intracortical inhibition (SICI) and facilitation (SICF) reflect a balance between the excitability of inhibitory and facilitatory circuits in the motor cortex.
SICI stimulus-response functions, measured with a range of conditioning stimulus intensities and two intervals (first peak and trough of the SICF function), were similar between younger and older adults, except at the shift from net inhibition to facilitation.
Relationships between SICI and SICF and manual dexterity show that an inhibition-facilitation balance favouring inhibition is associated with better manual dexterity and a balance favouring facilitation is associated with poorer manual dexterity.

**Abstract:** Research investigating age-related differences in intracortical inhibition acting within the primary motor cortex (M1) reports inconsistent results. Age-related changes in the balance of inhibition and facilitation acting within M1 might underlie the inconsistent findings. Here, paired-pulse TMS was used to examine the balance between short-interval intracortical inhibition (SICI) and short-interval intracortical facilitation (SICF) in younger (n=26) and older (n=21) adults. First, the inter-stimulus intervals (ISIs) that produced maximal and minimal SICF at the first peak and trough were identified for each individual. Second, SICI stimulus-response functions (conditioning stimulus (CS): 50–110% resting motor threshold) were determined at ISIs that produced maximal and minimal facilitation for each individual. When SICI was measured with the ISI that produced maximal facilitation, the shift from net inhibition to net facilitation occurred at a lower CS intensity in older than younger adults. When SICI was measured with the ISI that produced minimal facilitation, older adults showed a persistent net inhibition at the highest CS intensity, whereas younger adults showed a shift from net inhibition to net facilitation at high CS intensities. Relationships between SICI and SICF and manual dexterity were evident across all measures of the inhibition-facilitation balance, whereby a balance favouring inhibition is associated with better manual dexterity and a balance favouring facilitation is associated with poorer manual dexterity. These findings contribute to our understanding of age-related differences in intracortical inhibitory and facilitatory processes, as well as highlight the importance of considering the balance between inhibitory and facilitatory processes when measuring SICI and SICF.

## 1. Introduction

Aging is associated with a decline in manual dexterity, which can compromise the functional independence of older adults (Burke and Barnes 2006, Seidler et al. 2010). Growing evidence suggests that functional and structural changes in the primary motor cortex (M1) might, in part, underlie age-related decline in voluntary movement (Fling and Seidler 2012, Bhandari et al. 2016, Fujiyama et al. 2016). Both intracortical inhibitory and facilitatory processes acting within M1 are important for manual dexterity (Ziemann et al. 1996, Zoghi et al. 2003, Buccolieri et al. 2004, Cattaneo et al. 2005, Hammond and Vallence 2007, Reis et al. 2008, Cowie et al. 2016, Duque et al. 2017, Cirillo et al. 2018, Liao et al. 2018). However, the exact changes in these processes with advancing age are not well understood.

Paired-pulse transcranial magnetic stimulation (TMS) can be used to measure the excitability of short-interval intracortical facilitation (SICF) and short-interval intracortical inhibition (SICI). To measure SICF, a suprathreshold stimulus precedes a subthreshold stimulus by inter-stimulus intervals (ISI) of ~1.3–4.5 ms; the amplitude of the motor evoked potential (MEP) elicited by paired-pulse TMS is larger than the MEP elicited by a single suprathreshold TMS pulse. This facilitation of the MEP is observed at ISIs similar to the ~1.5 ms latency of indirect waves (I-waves; Ziemann et al. 1998, Di Lazzaro et al. 2012). SICF has three prominent peaks of facilitation: Peak 1 occurs at ISIs of 1.1—1.5 ms, Peak 2 occurs at ISIs of 2.3–2.9 ms, and Peak 3 occurs at ISIs of 4.1–4.4 ms (Tokimura et al. 1996, Ziemann et al. 1998, Di Lazzaro et al. 1999, Hanajima et al. 2002, Peurala et al. 2008). Between the three peaks are two troughs (ISIs: 1.6–2.2 ms and 3.0–4.0 ms), during which little or no facilitation occurs (Ziemann et al. 1998). The peaks of facilitation are thought to be the result of interactions between the excitatory postsynaptic potentials produced by the two stimuli within the intracortical facilitatory circuits that generate I-waves (Ziemann et al. 1998, Ziemann et al. 1998, Hanajima et al. 2002, Ortu et al. 2008, Di Lazzaro et al. 2012).

To measure SICI, a subthreshold conditioning stimulus (CS) precedes a suprathreshold test stimulus (TS) by ISIs of 1–5 ms; the amplitude of the MEP elicited by paired-pulse TMS is reduced compared to the MEP elicited by a single suprathreshold TMS pulse. This reduction in MEP amplitude is due to the activation of SICI circuits (Kujirai et al. 1993). Pharmacological studies provide strong evidence that SICI is mediated by GABA_A_ receptor activity (Ziemann et al. 1996, Ilic et al. 2002, Di Lazzaro et al. 2006). The contribution of the excitability of inhibitory interneurons that mediate SICI to the MEP can be measured by varying the CS intensity in the paired-pulse protocol: the paired-pulse MEP expressed as a ratio of the single-pulse MEP shows a U-shaped function with increasing CS intensity (Kujirai et al. 1993, Ziemann et al. 1996, Kossev et al. 2002, Orth et al. 2003). The descending limb of the SICI stimulus-response function, where SICI increases with increasing CS intensity, reflects the progressive recruitment of inhibitory interneurons that mediate SICI; the ascending limb, where SICI decreases with increasing CS intensity, reflects the progressive recruitment of excitatory interneurons.

A recent meta-analysis of studies investigating age-related changes in SICI using TMS showed inconsistent findings (Bhandari et al. 2016): some studies reported less SICI in older than younger adults (e.g., Peinemann et al. 2001, Marneweck et al. 2011), some reported no difference in SICI between older and younger adults (Oliviero et al. 2006, Rogasch et al. 2009, Cirillo et al. 2010, Cirillo et al., 2011, Fujiyama et al. 2011, Fujiyama et al. 2012, Opie and Semmler 2014), and some reported more SICI in older than younger adults (Smith et al. 2009, McGinley et al. 2010). The inconsistent SICI findings might be due to age-related changes in the balance between intracortical inhibitory and facilitatory processes acting within M1. Many studies measuring SICI in older adults have used a single CS intensity and/or a single ISI: although single CS intensities and single ISIs can be set to elicit net inhibition, this does not ensure an uncontaminated measure of SICI because activity in SICF circuits likely contributes to all SICI measures (albeit with varying magnitudes of SICF contribution (Di Lazzaro et al. 2008, Ni and Chen 2008, Rusu et al. 2014). Therefore, to understand age-related changes in SICI, it is important to consider the balance between SICI and SICF at a range of CS intensities and ISIs. Measures of SICI with varying contributions of SICF can be obtained by, first, identifying ISIs that evoke (i) maximal and (ii) minimal facilitation at the individual level and, second, measuring SICI at a range of CS intensities at the ISIs that produced maximal and minimal facilitation (Peurala et al. 2008). SICF will likely make a greater contribution to SICI measured at the first peak of maximal facilitation than SICI measured at minimal facilitation (i.e., the trough of the SICF function).

The current study investigated age-related differences in the balance between SICI and SICF. The first two SICF peaks and the trough between these peaks were obtained by varying the ISI from 1.3 ms to 3.1 ms (Peurala et al. 2008, Opie et al. 2018). Following the identification of ISIs that produced early maximal facilitation and minimal facilitation for each individual, SICI stimulus-response functions were obtained by varying CS intensity from 50–110% of resting motor threshold (10% increments). For each individual, two SICI stimulus-response functions were obtained: one using the individualised ISIs that produced maximal facilitation (SICF Peak 1) and one using the individualised ISIs that produced minimal facilitation (trough). Maximal facilitation was identified using ISIs that produced SICF Peak 1 because previous research shows that the magnitude of facilitation is greater at Peak 1 than Peak 2, and previous reports of age-related differences in SICF at Peak 1 are inconsistent (Clark et al. 2011, Opie et al. 2018). Finally, to explore the functional importance of SICI and SICF, correlation analyses were conducted between manual dexterity and (i) SICF at Peak 1 and Peak 2, and (ii) measures from the SICI stimulus-response functions at which net inhibition shifts to net facilitation.

## 2. Methods

### 2.1 Participants

Twenty-six younger adults (13 females; median 24 years, age range: 18–35 years) and 21 older adults (16 females; median 71 years, age range: 61–86 years) participated in this study. Participants were recruited from Murdoch University and the local community. The protocol was performed in accordance with the Declaration of Helsinki and approved by the Murdoch University Human Research Ethics Committee (2014–247). All participants gave written informed consent prior to testing and were screened for any conditions that would contraindicate TMS (Rossi et al. 2011, Rossini et al. 2015). The Montreal Cognitive Assessment was used to screen cognitive function. A cut-off score of 26 was used as an exclusion criterion; no participants received a score less than 26 (median 28, range 26–30; Nasreddine et al. 2005). The Edinburgh Handedness Inventory was used to measure handedness. A cut-off score of 40 was used as an exclusion criterion; no participants received a score less than 40 (Younger: median 100, score range: 75–100; Older: median 100, score range: 75–100; Oldfield 1971).

### 2.2 TMS

Participants were seated in a comfortable reclining chair with the right forearm supported by a cushion placed on their lap. Electromyographic (EMG) activity was recorded from the relaxed first dorsal interosseous (FDI) of the right hand using Ag-AgCI surface electrodes placed in a belly-tendon montage. The EMG signal was amplified (x1000; CED 1902 amplifier), bandpass filtered (20–1000 Hz) and digitized at a sampling rate of 2 kHz (CED 1401 interface). Two Magstim 200 stimulators connected via a BiStim module (Magstim Co., Whitland, Dyfed, UK) were used to generate (monophasic) single- and paired-pulses. Pulses were delivered through a figure-of-eight coil (90-mm diameter) placed tangentially to the left M1 with the handle positioned backwards and ~45° away from the midline to induce posterior-anterior current flow in the cortex. The optimal site, resting motor threshold (RMT), and TMS intensity to elicit MEPs ~1 mV in peak-to-peak amplitude (SI_1mV_) were obtained for the relaxed FDI muscle. The optimal site for stimulation was determined by systematically applying suprathreshold stimuli to identify a site that evoked consistent MEPs in the FDI. The optimal site for stimulation was marked on the scalp with water-soluble ink to allow for reliable coil placement throughout the experiment. RMT was defined as the minimum stimulus intensity (as a percentage of maximum stimulator output; MSO) required to elicit MEPs ≥ 50 μV in at least five out of 10 consecutive trials (Rossini et al. 2015). SI_1mV_ was defined as the stimulus intensity (as a % of MSO) required to evoke a MEP with a mean peak-to-peak amplitude of ~1 mV.

### 2.3 Experimental Protocol

The experiment took ~2.5 hours and followed the proceeding protocol: (1) manual dexterity was measured with the Purdue Pegboard test; (2) SICF was measured using single- and paired-pulse TMS; (3) individual ISIs that produced maximal and minimal SICF were determined for each subject; and (4) SICI stimulus-response functions were obtained at the individualised ISIs producing maximal and minimal facilitation using single- and paired-pulse TMS with varying CS intensities (order counterbalanced across participants).

#### Manual Dexterity

Manual dexterity was measured with the Purdue Pegboard unimanual sub-test, using the standardized testing procedure (Lafayette Instrument Company, USA). Participants were required to pick-up small pegs with their dominant right hand, one at a time from a well, and insert them from top to bottom into holes in a vertical array as quickly as possible. The number of pegs moved and placed in the 30 s period was measured.

#### Short-interval intracortical facilitation

SICF was obtained by delivering a suprathreshold stimulus (S1) at an intensity of SI_1mv_, followed by a subthreshold stimulus (S2) at an intensity of 90% RMT at 10 ISIs between 1.3 and 3.1 ms (steps of 0.2 ms; Peurala et al. 2008). Five experimental blocks, each consisting of 42 trials, were delivered. Each block consisted of 12 single-pulse (S1-alone) trials and three trials for each of the 10 paired-pulse conditions (i.e., 10 different ISIs). Trial conditions were pseudo-randomized with an inter-trial interval of 5 s (±20%). Each block lasted ~4 minutes and there was 3–5 minutes of rest between the five blocks.

To obtain individually optimised ISIs producing maximal and minimal SICF, the mean MEP amplitude was calculated for the single pulse condition and each of the 10 paired-pulse conditions. SICF was quantified by expressing the mean paired-pulse MEP amplitude for each ISI as a ratio of the mean single-pulse MEP amplitude. Ratios greater than 1.0 indicate facilitation. For each individual, the ISIs at which the SICF ratio was largest and smallest, showing maximal and minimal facilitation respectively, were subsequently used as the ISIs for the SICI stimulus-response functions (full details below).

#### Short-interval intracortical inhibition

SICI was obtained by delivering a subthreshold conditioning stimulus (CS) at seven intensities ranging from 50% to 110% RMT (steps of 10%), before a suprathreshold test stimulus (TS) at an intensity of SI_1mV_. Eight experimental blocks each consisting of 40 trials were performed: four blocks for the ISI producing maximal facilitation and four blocks for the ISI producing minimal facilitation (the order of the ISI tested first was counterbalanced across participants). Each block consisted of 12 single-pulse (TS-alone) trials and four trials for each of the seven paired-pulse conditions (i.e., seven different CS intensities). Trial conditions were pseudo-randomized with an inter-trial interval of 5 s (±20%). Each block lasted ~3.5 minutes and there was 3–5 minutes of rest between the four blocks.

### 2.4 Data Processing and Analysis

Trials in which peak-to-peak EMG activity exceeded 0.01 mV during the 100 ms prior to TMS application were excluded. The total % of trials excluded was 4.9% for younger adults and 5.3% for older adults. The peak-to-peak MEP amplitude was obtained from 40 ms of EMG activity beginning 10 ms after TMS application. Independent samples *t*-tests were performed to test for differences in RMT, SI_1mV_, and single-pulse MEP amplitude between younger and older adults.

#### SICF

In the current study, individual ISIs that produced SICF Peak 1 ranged between 1.3–1.5 ms and individual ISIs that produced SICF Peak 2 ranged between 2.5–3.1 ms. To test for differences in the magnitude of SICF peaks between younger and older adults, a mixed-factorial ANOVA was performed with the within-subject factor of ISI (Peak 1: 1.3, 1.5; Peak 2: 2.5, 2.7, 2.9, 3.1 ms) and between-subject factor of Age (younger, older).

#### SICI

To test for differences in SICI between younger and older adults, a mixed-factorial ANOVA was performed with the within-subject factor of CS intensity (50%, 60%, 70%, 80%, 90%, 100%, 110% RMT) and between-subject factor of Age (younger, older) using polynomial contrasts. Separate ANOVAs were performed for the SICI stimulus-response function obtained using the ISI producing maximal facilitation and the SICI stimulus-response function obtained using the ISI producing minimal facilitation.

Normality was violated for all mixed-factorial ANOVAs used to test for differences in SICF and SICI between younger and olders all; ratios were log-transformed before analysis and all analyses were performed on the log-transformed data. For ease of interpretation, nontransformed data are reported in the figures. Additionally, if the sphericity assumption was violated, Greenhouse-Geisser’s degrees of freedom adjustment was applied. All main effects and interactions were investigated using false-discovery-rate (FDR) post hoc tests (Curran-Everett 2000). All tests were two-tailed and statistical significance was accepted at an alpha level of *P* <.05. All values are expressed as mean and standard deviation (SD).

#### Association between manual dexterity and SICI and SICF

Pearson’s product moment correlation coefficients were obtained separately for younger and older adults to investigate the relationship between the manual dexterity (the number of pegs inserted on the Purdue Pegboard) and (i) SICF at Peak 1 and Peak 2, and (ii) measures from the SICI stimulus-response functions at CS intensities at which inhibition shifted to facilitation.

## 3 Results

### 3.1 Neurophysiological results

The mean RMT (% of MSO) was significantly higher in the older adult group (*M* = 59.1, *SD* = 9.2) than the younger adult group (*M* = 53.5, *SD* = 8.4; *t*_45_ = 2.17, *P* = 0.035, *d* = 0.64). Similarly, the mean SI_1mV_ (% of MSO) was significantly higher in the older adult group (*M* = 70.8, *SD* = 9.8) than the younger adult group (*M* = 61.9, *SD* = 10.2; *t*_45_ = 3.02, *P* = 0.004, *d* = 0.89). However, as expected, the mean amplitude of the single-pulse MEP evoked by SI_1mv_ was similar in younger (*M* = 1.0 mV, *SD* = 0.5) and older (*M* = 0.9 mV, *SD* = 0.9) adult groups (*t*_45_ = 0.89, *P* = 0.378, *d* = 0.26).

#### SICF

Figure 1 shows SICF as a function of ISI in younger and older adults. Both groups showed two similar peaks of facilitation (Peak 1: 1.3–1.5 ms; Peak 2: 2.5–3.1 ms) separated by a trough (1.9–2.3 ms). Both the group (Fig 1A) and individual data (Fig 1B and 1C) show a slower shift from the trough to the second peak of facilitation in older than younger adults. An ANOVA performed on the SICF data at the two peaks showed no main effect of Age (*F*_1_, 45 = 0.100, *P* = 0.753, η_p_^2^ = 0.00), but a significant main effect of ISI (*F*_2.94, 132.30_ = 14.95, *P* < 0.001, η_p_^2^ = 0.25) and a significant Age*ISI interaction (*F*_2.94, 132.30_ = 2.81, *P* = 0.043, η_p_^2^ = 0.06). FDR calculations were performed to examine the difference in SICF between younger and older adults. Consistent with previous research, the younger adult group showed significantly greater facilitation than the older adult group at the 2.5 ms ISI (*t*_45_ = 2.17, *P* = 0.035, *d* = 0.63). One younger participant shows extreme facilitation (MEP ratio greater than 4.0): when this extreme facilitator was removed from the analysis, younger adults still showed significantly greater facilitation than older adults at the 2.5 ms ISI: *t*_44_ = 2.08, *P* = 0.046, *d* = 0.56). There was no significant difference in SICF between younger and older adults at any of the other ISIs (all *t*_45_ < 1.21, all *P* > 0.234, all *d* < 0.32).

**Figure 1.**
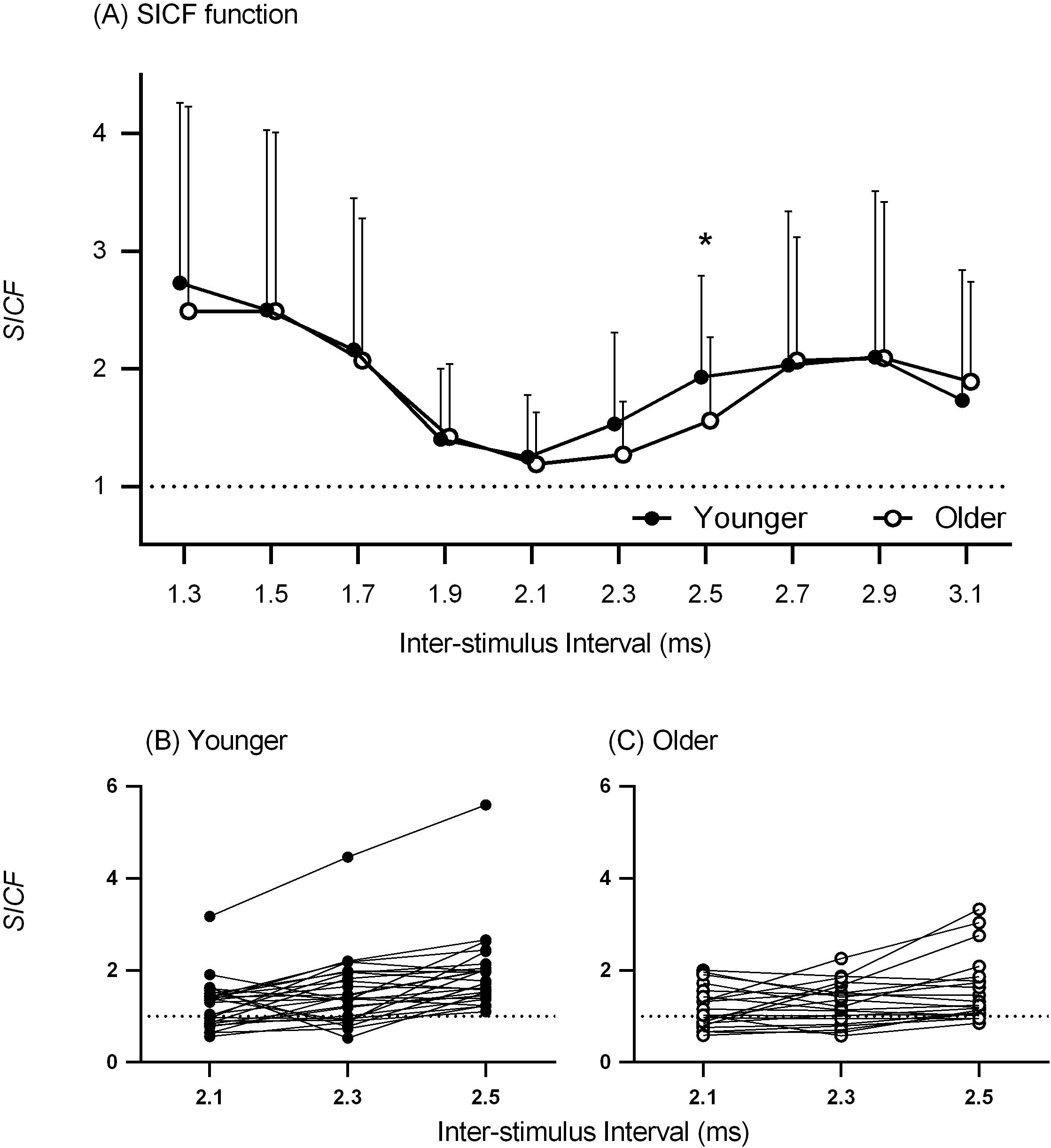
Panel A shows SICF (mean paired-pulse MEP amplitude expressed as a ratio of mean single-pulse MEP amplitude) for each of the 10 ISIs in younger (closed symbols) and older adults (open symbols). A ratio greater than 1.0 reflects facilitation. The error bars show the upper limit standard deviation. Panels B and C show individual data at ISIs of 2.1 ms, 2.3 ms, and 2.5 ms for younger and older adults, respectively. * difference between younger and older, *P* <.05.

#### SICI

To obtain measures of SICI with varying contributions of SICF, the ISIs that produced maximal and minimal facilitation for each individual were used as the ISIs in the SICI stimulus-response function protocol. SICF will likely make a greater contribution to SICI measured at the ISI producing maximal facilitation than SICI measured at the ISI producing minimal facilitation. Maximal facilitation occurred at ISIs of 1.3 ms and 1.5 ms in both younger and older adults. Minimal facilitation ranged across a larger number of ISIs than maximal facilitation in younger (1.7–2.3 ms) and older adults (1.9–2.3 ms). To measure SICI with varying contributions of SICF, SICI stimulus-response functions were obtained using a range of CS intensities. We report below SICI stimulus-response functions first, at the ISI that produced maximal facilitation and second, at the ISI that produced minimal facilitation.

#### SICI measured at the ISI producing maximal facilitation

Figure 2 shows SICI stimulus-response functions measured at the ISI producing maximal facilitation in younger and older adults. The pattern of SICI with increasing CS intensity is similar in younger and older adults; specifically, a progressive increase in inhibition with CS intensity at the lower CS intensities (50–70% RMT), followed by a shift from inhibition to facilitation at higher CS intensities (80–110% RMT). However, there is less inhibition (i.e., larger ratios) at the CS intensity of 80% RMT in older than younger adults (individual data shown in Fig 2B and 2C), indicating a shift from net inhibition to net facilitation at a lower CS intensity in older than younger adults. An ANOVA performed on the SICI data measured at the ISI producing maximal facilitation showed no main effect of Age (*F*_1, 45_ = 1.68, *P* = 0.202, η_p_^2^ = 0.04), but a significant main effect of CS intensity (*F*_1, 45_ = 86.20, *P* < 0.001, η_p_^2^ = 0.66) and a significant Age*CS intensity interaction (*F*_1, 45_ = 9.28, *P* = 0.004, η_p_^2^ = 0.17). FDR calculations were performed to examine the difference in SICI between younger and older adults at each of the CS intensities. At the CS intensity of 80% RMT, the older adult group showed significantly less inhibition than the younger adult group (*t*_45_ = 2.69, *P* = 0.010, two-tailed, *d* = 0.80). There was no significant difference in SICI between younger and older adults at any of the other CS intensities (all *t*_45_ < 1.81, all *P* > 0.077, all *d* < 0.54; the CS intensity 70% RMT showed a medium effect size, with the older adult group showing less inhibition than the younger adult group).

**Figure 2.**
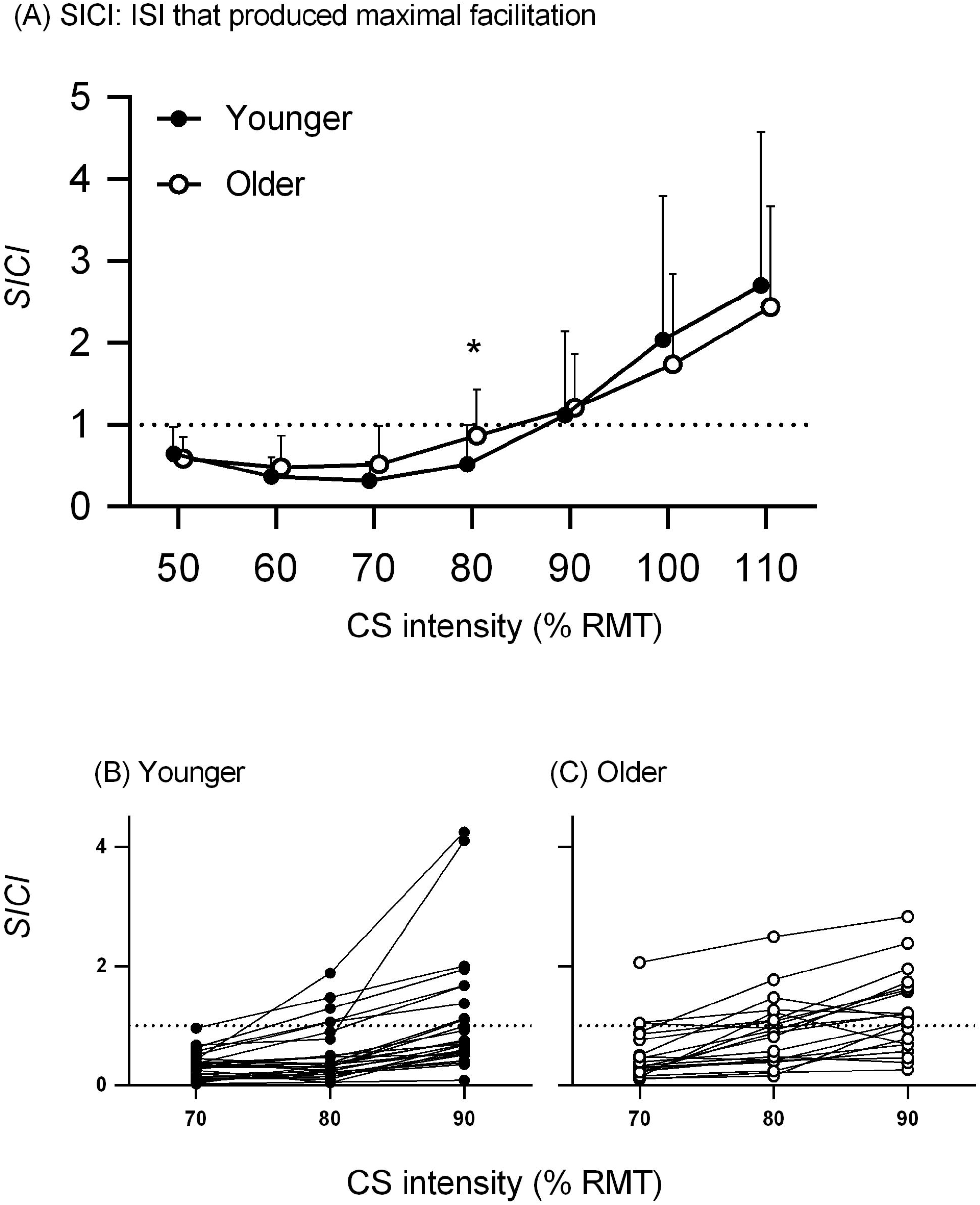
Panel (A) shows SICI (mean paired-pulse MEP amplitude expressed as a ratio of mean single-pulse MEP amplitude) for each of the seven CS intensities in younger (closed symbols) and older adults (open symbols) measured at the ISI producing maximal facilitation (SICF Peak 1). A ratio less than 1.0 reflects inhibition; smaller ratios reflect greater inhibition. The error bars show the upper limit standard deviation. Panels B and C show individual data at CS intensities 70–90% RMT for younger and older adults, respectively. * difference between younger and older, *P* <.05.

#### SICI measured at the ISI producing minimal facilitation

Figure 3 shows SICI stimulus-response functions measured at the ISI producing minimal facilitation in younger and older adults. It can be seen from Figure 3A that the younger adult group showed the expected U-shaped stimulus-response function with increasing CS intensity, but the older adult group showed a flatter function, with very little change in SICI across the range of CS intensities tested. The data show more inhibition (i.e., smaller ratios) at the highest CS intensities (100–110% RMT) in older than younger adults (Fig 3B and 3C); few older adults showed ratios greater than 1.0, indicating a shift from net inhibition to net facilitation in younger but not older adults. An ANOVA performed on the SICI data measured at the ISI that produced minimal facilitation showed no main effect of Age (*F*_1, 45_ = 0.58, *P* = 0.450, η_p_^2^ = 0.01), but a significant main effect of CS intensity (*F*_1, 45_ = 63.55, *P* < 0.001, η_p_^2^ = 0.59) and a significant Age*CS intensity interaction (*F*_1, 45_ = 7.72, *P* = 0.008, η_p_^2^ = 0.15). FDR calculations were performed to examine the difference in SICI between younger and older adults at each of the CS intensities. At both the lowest and the highest CS intensities, the younger adult group showed less inhibition than the older adults group. Specifically, at the CS intensity of 50% RMT, the younger adult group showed significantly less inhibition than the older adult group (*t*_45_ = 2.84, *P* = 0.007, *d* = 0.82); at the CS intensity of 60% RMT, the younger adult group also showed less inhibition than the older adult group but this difference failed to reach statistical significance (*t*_45_ = 2.00, *P* = 0.051, *d* = 0.60). At the CS intensity of 110% RMT, the younger adult group show significantly larger ratios, indicating less inhibition or a shift to facilitation, than the older adult group (*t_43.57_* = 2.59, *P* = 0.013, *d* = 0.74). There was no significant difference in SICI between younger and older adults at any of the other CS intensities (all *t*_45_ < 1.16, all *P* > 0.252, *d* < 0.52).

**Figure 3.**
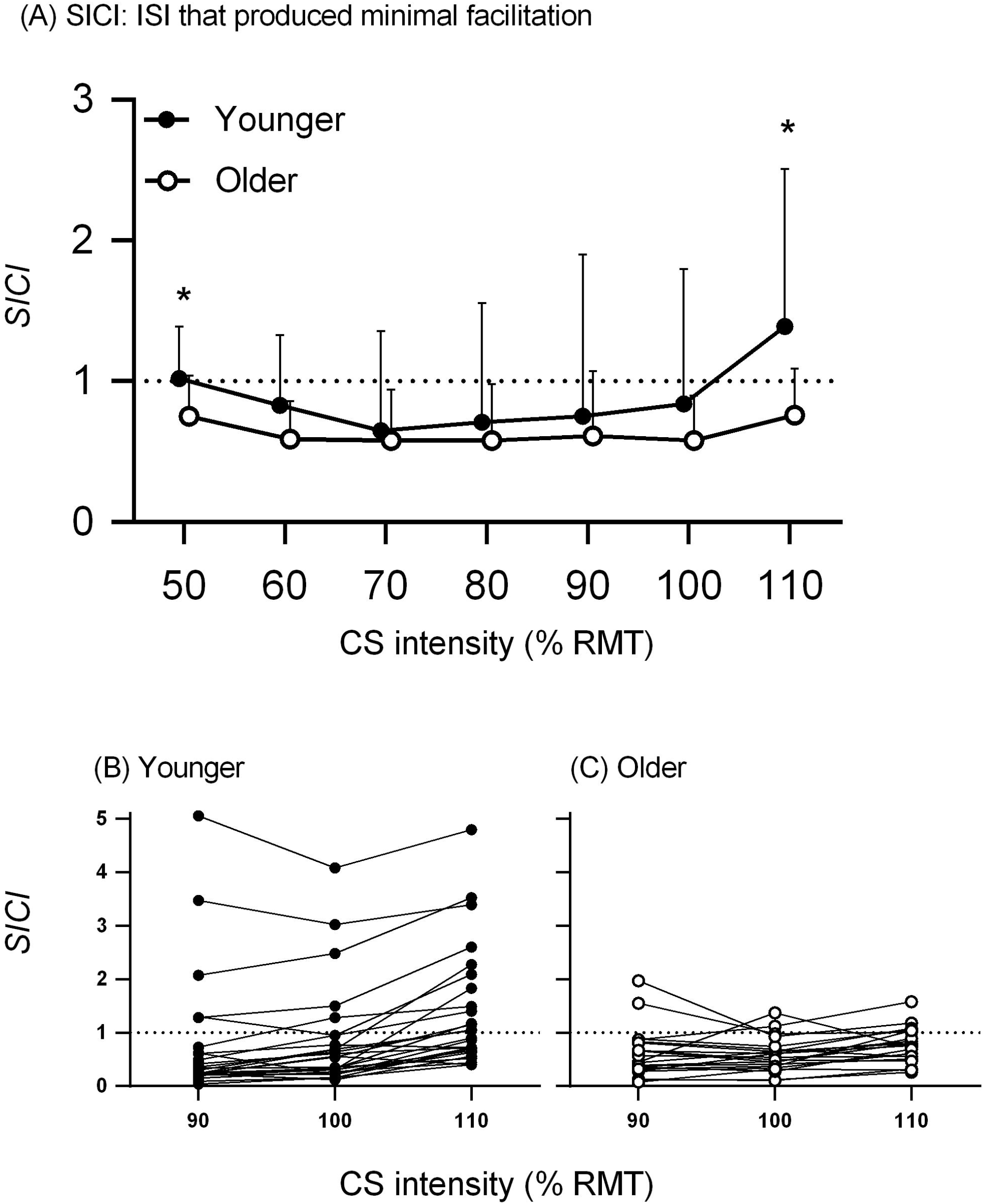
Panel (A) shows SICI ratios (mean paired-pulse MEP amplitude expressed as a ratio of mean single-pulse MEP amplitude) for each of the seven CS intensities in younger (closed symbols) and older adults (open symbols) measured at the ISI producing minimal facilitation (SICF Trough). A ratio less than 1.0 reflects inhibition; smaller ratios reflect greater inhibition. The error bars show the upper limit standard deviation. Panels B and C show individual data at CS intensities 90–110% RMT for younger and older adults, respectively. * difference between younger and older, *P* <.05.

#### Associations between manual dexterity and inhibition-facilitation balance

On the unimanual Purdue Pegboard subtest, older adults placed significantly fewer pegs (*M* = 12.4, *SD* = 2.5) than younger adults (*M*= 15.6, *SD* = 1.9; independent samples t-test: *t*_45_ = 5.04, *P* < 0.001, *d* = 1.48). To explore the functional importance of SICF and SICI in manual dexterity, correlation analyses were performed between Purdue pegboard performance and (i) SICF Peak 1 and 2, and (ii) SICI at the CS intensities of 80% RMT and 110% RMT at the ISIs that produced maximal and minimal facilitation. CS intensities 80% RMT and 110% RMT were selected because they were the intensities at which net inhibition differed between younger and older adults at the ISIs that produced maximal and minimal facilitation, respectively, and likely reflect the shift from net inhibition to net facilitation. Figure 4 shows scatterplots of the relationship between number of pegs placed and SICF at Peak 1 and Peak 2. In younger adults, a significant moderate negative correlation was found between the number of pegs placed and SICF at Peak 1 (based on the individual ISI that produced Peak 1 (range 1.3–1.5 ms): r26 = −0.45, *P* = 0.021), showing that more facilitation was associated with poorer Purdue Pegboard performance (i.e., less pegs placed). The scatterplot shows four individuals with extreme facilitation (i.e., ratios > 4.0): when the extreme facilitators were removed from the analysis, the correlation between SICF at Peak 1 and Purdue Pegboard performance was smaller than the correlation with the full sample (*r*_22_ = −0.20, *P* = 0.368). In younger adults, there was also a significant moderate negative correlation found between the number of pegs placed and SICF at Peak 2 (based on the individual ISI that produced Peak 2 (range 2.5–3.1 ms): *r*_26_ = −0.51, *P* = 0.008), also showing that more facilitation was associated with poorer Purdue Pegboard performance. The scatterplot shows one individual with extreme facilitation (i.e., ratio > 4.0): when the extreme facilitator was removed from the analysis, the correlation between SICF at Peak 2 and Purdue Pegboard performance was similar to the full sample (*r*_25_ = −0.57, *P* = 0.003). In older adults, there were no significant correlations between Purdue Pegboard performance and SICF at either Peak 1 or Peak 2 (both *r*_21_ < −0.25, all *P* > 0.270); correlations remained small and non-significant when the one older adult extreme facilitator was removed from the analyses (both *r*_20_ < −0.14, all *P* > 0.570).

**Figure 4.**
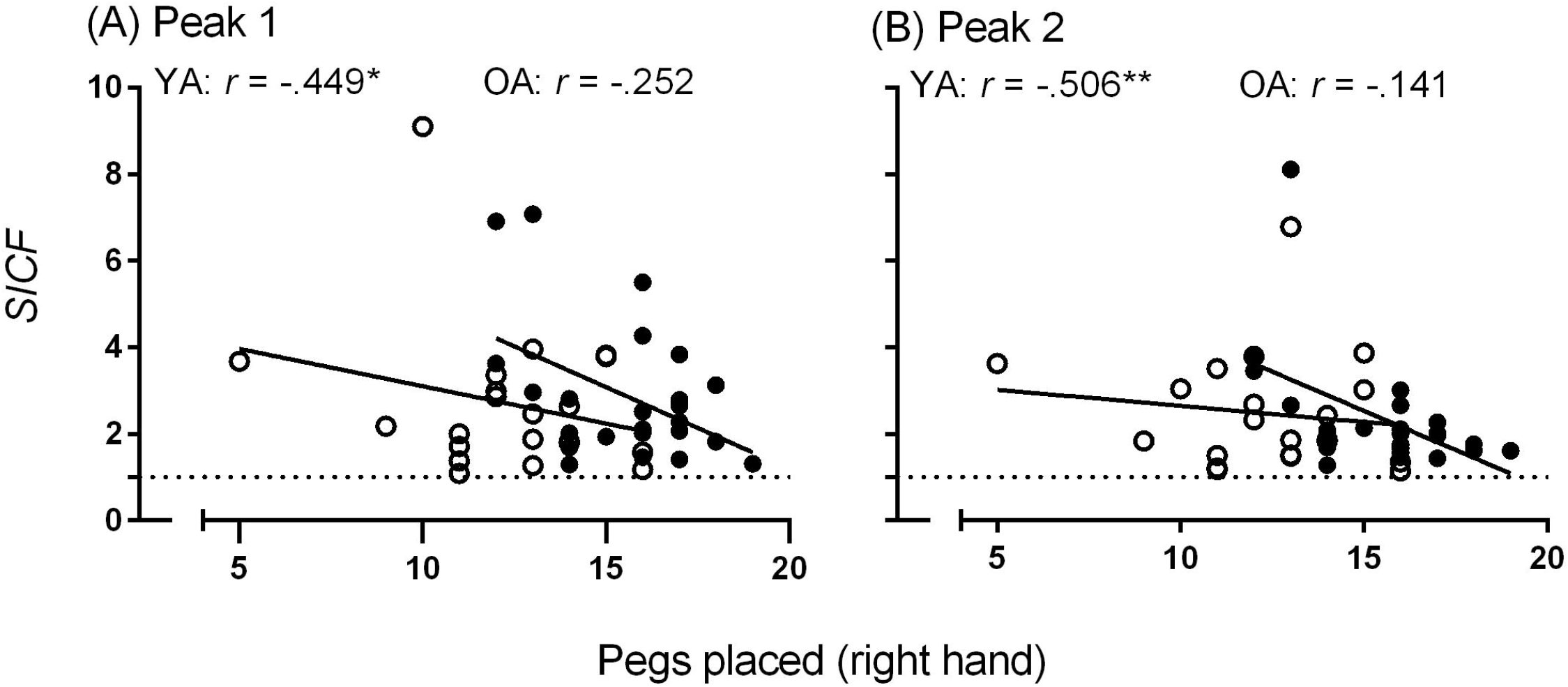
Scatterplots of the relationship between the performance on the Purdue Pegboard and the magnitude of SICF at individual ISIs that produced Peak 1 (A) and Peak 2 (B) for younger (closed symbols) and older adults (open symbols). Lines of best fit are represented as a solid line for younger and dotted line for older adults. \**P* <.05; ** *P* <.01

Figure 5 shows the scatterplots of the relationship between the performance on the Purdue Pegboard and SICI measured at the ISI producing maximal facilitation (top) and minimal facilitation (bottom). In younger adults, for SICI measured at the ISI that produced maximal facilitation (top), a moderate significant negative correlation was found between Purdue Pegboard performance and SICI at CS intensity 110% RMT (*r*_26_ = −0.52, *P* = 0.007), showing that more inhibition (or less facilitation) was associated with more pegs inserted in the Purdue Pegboard subtest. This is consistent with the association between SICF at Peak 1 and Purdue performance reported above. In younger adults, there were no significant correlations found between Purdue Pegboard performance and SICI at the CS intensity 80% RMT (*r*_26_ = −0.12, *P* = 0.560). In older adults, for SICI measured at the ISI producing maximal facilitation, there were no significant correlations between Purdue pegboard performance and SICI at either of the CS intensities (*r*_21_ = −0.08, *P* = 0.726).

**Figure 5.**
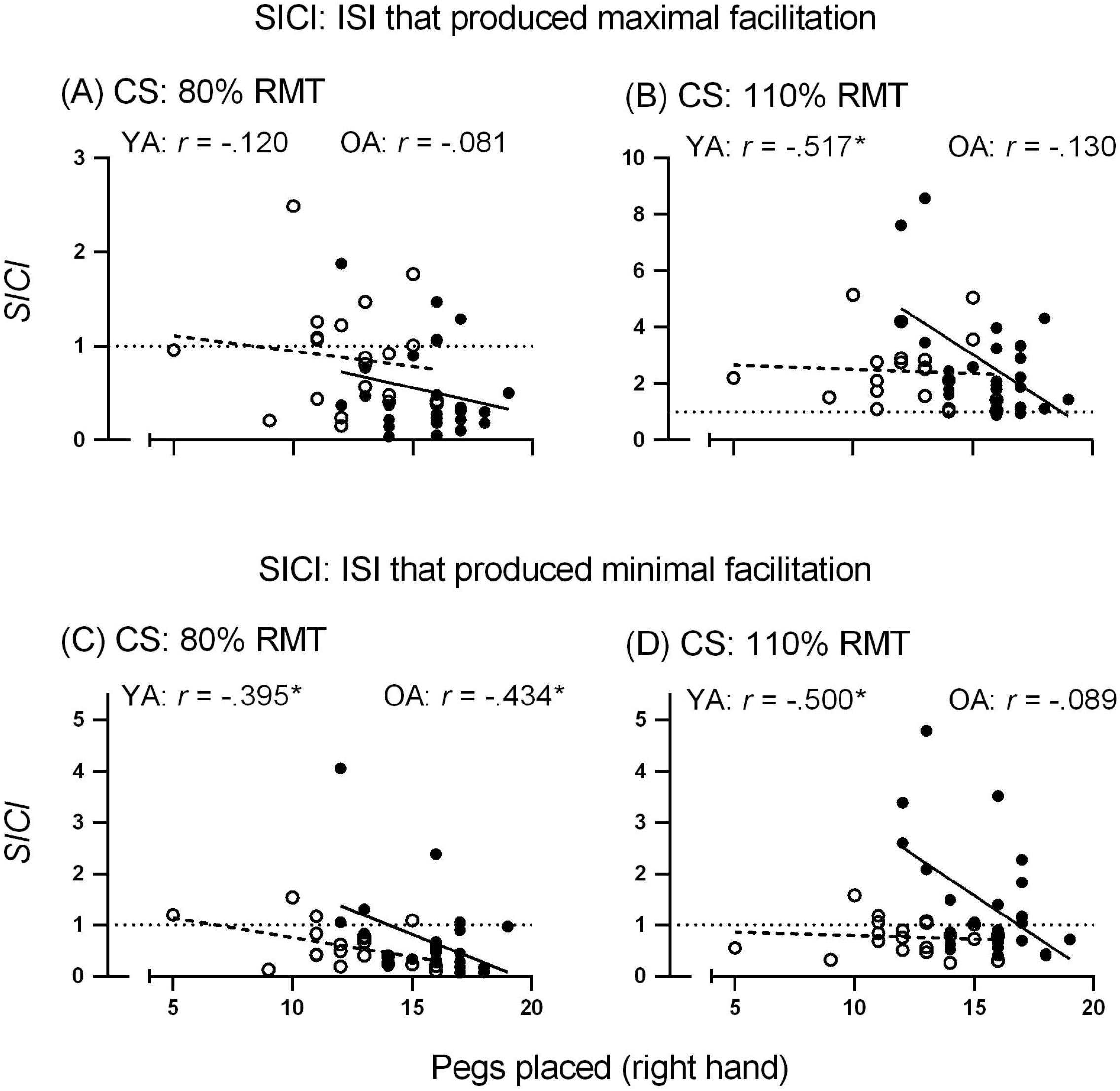
Scatterplots of the relationship between the performance on the Purdue Pegboard and the magnitude of SICI measured at the ISI producing maximal facilitation (top row) and minimal facilitation (bottom row) for younger (closed symbols) and older adults (open symbols). Panels (A) and (C) show SICI ratios at the CS intensity of 80%RMT and panels (B) and (D) show SICI ratios at the CS intensity of 110% RMT. Lines of best fit are represented as a solid line for younger and dotted line for older adults. \**P* <.05.

For SICI measured at the ISI producing minimal facilitation (Fig 5; bottom), in younger adults, significant moderate negative correlations were found between Purdue Pegboard performance and SICI at CS intensities of 80% RMT (*r*_26_ = −0.40, *P* = 0.046) and 110% RMT (*r*_26_ = −0.50, *P* = 0.009). These correlations show that more inhibition is associated with more pegs inserted (i.e., better performance). In older adults, a significant moderate negative correlation was found between Purdue Pegboard performance and SICI at the CS intensity 80% RMT (*r*_21_ = −0.43, *P* = 0.048), showing that more inhibition is associated with more pegs inserted. In older adults, there was no significant correlation between Purdue Pegboard performance and SICI at the CS intensity 110% RMT (*r*_21_ = −0.09, *P* = 0.702).

## 4. Discussion

The current study characterised the balance between the excitability of short-acting facilitatory and inhibitory circuits across the 1.3–3.1 ms time course in younger and older adults. At ISIs that produced maximal and minimal facilitation, stimulus-response functions showed that the shift from net inhibition to net facilitation differed between younger and older adult groups. When SICI was measured with the ISI that produced maximal facilitation (1.3–1.5 ms), the reduction in inhibition that preceded the shift from net inhibition to net facilitation occurred at a lower CS intensity in the older adult group than the younger adult group (Fig 2). When SICI was measured with the ISI that produced minimal facilitation (1.7–2.3 ms), older adults showed a persistent net inhibition at the highest CS intensity, whereas younger adults showed a shift from net inhibition to net facilitation at high CS intensities (Fig 3); this persistent net inhibition at the highest CS intensity in older but not younger adults likely contributed to the onset of maximal facilitation at SICF Peak 2 occurring at longer ISIs in older than younger adults and the reduced SICF Peak 2 at the 2.5 ms ISI in older than younger adults. Finally, the current findings provide some evidence for the relationship between inhibitory-facilitation balance and manual dexterity, whereby a balance favouring inhibition is associated with better manual dexterity and a balance favouring facilitation is associated with poorer manual dexterity.

### Age-related differences in SICF

The magnitude of facilitation at the ISIs that produced the first peak of the SICF function was similar in younger and older adult groups, suggesting that the excitability of the summed excitatory synaptic activity within the intracortical facilitatory circuits that generate the first I-wave is similar in younger and older adults (Ziemann et al. 1998, Ziemann et al. 1998, Hanajima et al. 2002, Ortu et al. 2008, Di Lazzaro et al. 2012). Reports of age-related differences in SICF Peak 1 are inconsistent: Clark et al. (2011) showed greater SICF (at an ISI of 1.5 ms) in older than younger adults, and Opie et al. (2018) showed smaller SICF (at both 1.3 ms and 1.5 ms) in older than younger adults. The two previous studies and the current study used the same stimulation intensities for single- and paired-pulse trials, and at least a subset of the same ISIs. Although the statistical outcomes of the three studies investigating SICF in younger and older adults are conflicting, the magnitude of facilitation at Peak 1 (~250%) for younger adults is similar across the three studies. It remains unclear why the three studies show inconsistent age-related differences in SICF Peak 1. At the ISI of 2.5 ms, the current study and the two previous studies showed reduced SICF in older than younger adults (Clark et al. 2011, Opie et al. 2018). In the current study, maximal facilitation at the second peak of the SICF function occurred at a longer ISI in the older adult group than the younger adult group, which is consistent with the results of Opie et al. (2018) that also showed maximal SICF Peak 2 at longer ISIs in older than younger adults. This temporal shift in the onset of maximal SICF Peak 2 in older adults can explain (in part) the smaller SICF at the ISI of 2.5 ms in older than younger adults, which has been reported consistently (Clark et al. 2011, Opie 2018). Taken together, these findings suggest that the maximal summed excitatory synaptic activity within the intracortical facilitatory circuits that generate late I waves occurs later in older than younger adults (Ziemann et al. 1998, Ziemann et al. 1998, Hanajima et al. 2002, Ortu et al. 2008, Di Lazzaro et al. 2012). Evidence suggests that different mechanisms contribute to SICF at early (Peak 1) and later (Peaks 2 and 3) peaks (Wagle-Shukla et al. 2009, Shirota et al. 2010, Cirillo et al. 2016): when taken together with the current SICF Peak 1 results, the SICF Peak 2 results suggest that advancing age slows the activity within circuits generating I-waves that contribute to the later peaks, but not the early peak of SICF. Opie et al. (2018) suggest that the longer latency of SICF Peak 2 in older than younger adults is unlikely to be due to the slowing of conduction velocity in the corticospinal tract, but likely due to the influence of intracortical inhibition on the later peaks of facilitation.

### Age-related differences in SICI measured at the ISI that produced maximal facilitation

Early in the SICI-SICF time course, at the ISIs that produced maximal facilitation (1.3–1.5 ms), the shift from net inhibition to net facilitation in the SICI stimulus-response function began at a lower CS intensity in older than younger adults. At these short ISIs, the SICI stimulus-response function likely reflects an early phase of inhibition present at an ISI of ~1 ms (Fisher et al. 2002, Hanajima et al. 2003, Roshan et al. 2003). The mechanisms underlying the early phase of SICI are still debated. In part, the early phase of SICI likely reflects axonal refractoriness; however, evidence suggests that extrasynaptic GABA_A_ activity and synaptic inhibition, mediated by low-threshold inhibitory interneurons, also contributes to the early phase of SICI (Fisher et al. 2002, Hanajima et al. 2003, Roshan et al. 2003, Ni et al. 2007, Vucic et al. 2009). If the early phase of SICI reflected only neuronal refractoriness, higher CS intensities would activate a larger pool of interneurons leading to greater inhibition: our findings showing the shift from inhibition to facilitation at higher CS intensities (90–110% RMT), suggesting that refractoriness of excitatory interneurons cannot fully explain the early phase of SICI. Previous studies that investigated age differences in SICI at an ISI of 1 ms showed less SICI in older than younger adults with a CS intensity of ~80% RMT (Peinemann et al. 2001, Smith et al. 2009), and Smith et al. (2009) showed a trend towards a reduction in inhibition (and shift towards facilitation) at lower CS intensities in older than younger adults (Smith et al. 2009), which is consistent with the current results. The shift from net inhibition to net facilitation at a lower CS intensity in older adults than younger adults suggests that activity within the SICF circuits that generate the early I-wave overrides the effect of activity within SICI circuits that mediate the early phase of inhibition at a lower intensity in older than younger adults. Given that there was no difference in the magnitude or latency of SICF Peak 1 between younger and older adult groups in the current study, these findings suggest that the efficacy of the early phase of SICI is reduced in older than younger adults.

### Age-related differences in SICI measured at the ISI that produced minimal facilitation

Later in the SICI-SICF time course, at the ISI that produced minimal facilitation (1.7–2.3 ms), younger adults showed a shallow U-shaped stimulus response curve, with a shift from net inhibition to net facilitation at high CS intensities, whereas older adults showed a persistent net inhibition at all CS intensities tested. At these ISIs, the SICI stimulus-response function likely reflects the activity of a second, later phase of GABA_A_ receptor activity mediated inhibition present at ISIs ≥ 2 ms (Kujirai et al. 1993, Ziemann et al. 1996, Ziemann et al. 1996, Ilic et al. 2002, Vucic et al. 2009). Two previous studies have examined age-related differences SICI using a 2 ms ISI, which is comparable to the ISIs that produced minimal facilitation in the current study (1.7–2.3 ms): one study reported less SICI in older than younger adults (Marneweck et al. 2011) and another study reported no age difference in SICI (Opie et al. 2015) (both studies used a single CS intensity: ~70%RMT). The current study is consistent with the report of Opie et al. (2015) showing no age difference in SICI at moderate intensities, and extends on this work to show a persistent net inhibition at high CS intensities in older but not younger adults. Taken together, these findings suggest increased strength but reduced sensitivity (to changing stimulus input) of SICI circuits mediated by GABA_A_ receptor activity in older than younger adults. Furthermore, this persistent and strong inhibition at high CS intensities in older adults likely contributes to the later onset of maximal facilitation at SICF Peak 2 and the reduced magnitude of SICF Peak 2 at the ISI of 2.5 ms in older compared to younger adults.

The second phase of GABA_A_ receptor activity mediated inhibition is also reflected in SICI measured at a 3 ms ISI. Most studies measuring SICI at an ISI of 3 ms and moderate CS intensities (~60–80% RMT) reported no difference between younger and older adults (Rogasch et al. 2009, Smith et al. 2009, Cirillo et al. 2010, Cirillo et al. 2011, Fujiyama et al. 2011, Fujiyama et al. 2012, Opie and Semmler 2014), although there is one report of reduced SICI in older than younger adults (Peinemann et al. 2001). At high CS intensities (~90–100% RMT) and an ISI of 3 ms, it has been reported that older adults show less inhibition than younger adults (Smith et al. 2009), potentially reflecting a shift from net inhibition to net facilitation at lower CS intensities in older than younger adults. Although the mean ratio for older adults was close to 1.0, indicating neither net inhibition nor net facilitation at the group level at the highest intensity, it is plausible that activity within the SICF circuits that generate the later I-wave overrides the effect of activity within SICI circuits that mediate the later phase of inhibition at a lower intensity in older than younger adults. To better understand the association between SICI circuits and SICF Peak 2, future research is required to obtain a SICI stimulus-response function at the ISI that produced maximal facilitation over the time course of SICF Peak 2.

### Relationship between manual dexterity and SICF and SICI

Insight into the associations between manual dexterity and the inhibition-facilitation balance that favours facilitation can be obtained from measures of (i) SICF and (ii) SICI measured at the ISI that produced maximal facilitation and a high CS intensity (110% RMT). In younger adults, moderate negative associations were found between Purdue Pegboard performance and these measures of the inhibition-facilitation balance, showing that greater facilitation was associated with fewer pegs placed. It is clear from the scatterplots (Fig 4A and 4B and Fig 5B) that younger adults who show an inhibition-facilitation balance strongly favouring facilitation show the poorest Purdue Pegboard performance. This is consistent with previous research showing that individuals with atypical facilitation—that is, net facilitation in response to a paired-pulse protocol set to measure SICI—performed more poorly on the Purdue pegboard task than individuals with net inhibition (Marneweck et al., 2011). The results suggest that, in younger adults, strong facilitation is associated with poor performance of movements of the fingers, hand, and arm required for successful completion of the Purdue Pegboard. In older adults, there was no association between performance in Purdue Pegboard task and either (i) SICF Peak 1 or (ii) SICI measured at the ISI that produced maximal facilitation at the high CS intensity 110% RMT. It is not clear why SICF at Peak 1 is associated with manual dexterity in younger but not older adults, but it might be due in part to age-related changes in both cortical processes and peripheral factors, such as cutaneous sensory perception (Decorps et al. 2014), proprioception (Goble et al. 2009) and muscle strength (Sosnoff and Newell 2006). Finally, in both younger and older adults, there were no associations between SICI measured at the ISI that produced maximal facilitation at the moderate CS intensity 80% RMT, suggesting less involvement of early phase of SICI and manual dexterity.

Insight into the associations between manual dexterity and the inhibition-facilitation balance that favours inhibition can be obtained from SICI measured at the ISI that produced minimal facilitation and a moderate CS intensity (80% RMT). In both younger and older adults, associations were found between Purdue Pegboard performance and this measure of SICI, showing greater inhibition was associated with more pegs placed. This positive association between Purdue performance and an inhibition-facilitation balance that favours inhibition suggests that the second phase of SICI, which is mediated by GABA_A_ receptor activity, is important for manual dexterity in both younger and older adults, potentially due to a role of GABAergic intracortical inhibitory circuits controlling neuronal firing in M1 (Farrant & Nusser, 2005; Heise et al., 2013).

When SICI was measured at the ISI that produced minimal facilitation at the high CS intensity of 110% RMT, younger adults showed net facilitation (at the group level) and there was a significant association with Purdue Pegboard performance, showing that greater facilitation was associated with fewer pegs placed. This finding provides further support for the suggestion that an inhibition-facilitation balance that strongly favours facilitation is associated with poor Purdue Pegboard performance in younger adults. In contrast, older adults showed net inhibition (at the group level) when SICI was measured at the high CS intensity, and there was no association with Purdue Pegboard performance. At first glance, this finding suggests that the persistent inhibition evident at higher CS intensities (reflecting the second phase of SICI), in older adults is not functionally beneficial, which fits with a previous suggestion that persistent inhibition in older adults might negatively affect the synchronization of neuronal firing in M1 and result in slowed motor processing (Heise et al., 2013). However, when taken together with the negative association between an inhibition-facilitation balance favouring facilitation and manual dexterity observed in younger adults, it is possible that the persistent inhibition in older adults might be a compensatory mechanism to moderate the negative effect of strong facilitation on manual dexterity, albeit at the expense of movement speed.

### Conclusions

The current study characterised the balance between excitability of the SICF circuits that generate I-waves and the GABAergic circuits that mediate SICI in younger and older adults. Early in the SICI-SICF time course (~1.5 ms), the shift from net inhibition to net facilitation occurred at lower CS intensities in older adults than younger adults, suggesting reduced efficacy of the early phase of SICI in older than younger adults. Later in the SICI-SICF time course (~2 ms), younger adults showed a shift to net facilitation at high CS intensities whereas older adults showed a persistent net inhibition, which likely contributes to the reduced SICF Peak 2 (at 2.5 ms) in older than younger adults that is consistent in the literature. Finally, evidence for relationships between SICI and SICF and manual dexterity were evident across all measures of the inhibition-facilitation balance, whereby a balance favouring inhibition is associated with better manual dexterity and a balance favouring facilitation is associated with poorer manual dexterity.

## Abbreviations

CS: conditioning stimulus;
EMG: electromyography;
FDI: first dorsal interosseous;
ISI: interstimulus interval;
M1: primary motor cortex;
MEP: motor evoked potential;
MSO: maximum stimulus output;
RMT: resting motor threshold;
SICF: short-interval intracortical facilitation;
SICI: short-interval intracortical inhibition;
TMS: transcranial magnetic stimulation.

